# The integrated stress response/eIF2a pathway controls cytokine production in tissue-resident memory CD4^+^ T cells

**DOI:** 10.1101/2024.01.26.577246

**Authors:** Nariaki Asada, Pauline Ginsberg, Hans-Joachim Paust, Ning Song, Jan-Hendrik Riedel, Jan-Eric Turner, Anett Peters, Anna Kaffke, Jonas Engeßer, Huiying Wang, Yu Zhao, Philipp Gild, Roland Dahlem, Sarada Das, Zoya Ignatova, Tobias B. Huber, Immo Prinz, Nicola Gagliani, Hans-Willi Mittrücker, Christian F. Krebs, Ulf Panzer

**Affiliations:** III. Department of Medicine, University Medical Center Hamburg-Eppendorf, Germany; Hamburg Center for Translational Immunology, University Medical Center Hamburg-Eppendorf, Germany; Institute of Medical Systems Biology, Center for Biomedical AI, Center for Molecular Neurobiology Hamburg, Germany; Department of Urology, University Medical Center Hamburg-Eppendorf, Germany; Institute of Biochemistry and Molecular Biology, University of Hamburg, Germany; Hamburg Center for Kidney Health (HCKH), University Medical Center Hamburg-Eppendorf, Germany; Institute of Systems Immunology, University Medical Center Hamburg-Eppendorf, Germany; Department of General, Visceral and Thoracic Surgery, University Medical Center Hamburg-Eppendorf, Germany; I. Department of Medicine, University Medical Center Hamburg-Eppendorf, Germany; Institute for Immunology, University Medical Center Hamburg-Eppendorf, Germany

## Abstract

Tissue-resident memory T (Trm) cells are a specialized T cell population that resides in tissues and can play both a protective and pathogenic role. The mechanism that enables Trm cells to provide a rapid protective response while restricting their function in homeostasis remains unclear. Here, we show that human and mouse CD4^+^ Trm cells exist in a *poised* state, characterized by storage of proinflammatory type-1 and type-3 cytokine mRNAs without protein production. In steady-state conditions, cytokine mRNA translation in Trm cells is suppressed by the integrated stress response (ISR)/eIF2α pathway, whereas Trm-cell activation under inflammatory conditions results in eIF2α dephosphorylation, leading to derepression and rapid translation of the cytokine mRNAs stored in stress granules. Pharmacological inhibition of eIF2α dephosphorylation resulted in reduced cytokine production from Trm cells, and ameliorated autoimmune kidney disease in mice. Consistent with these results, the ISR pathway in Trm cells was downregulated in patients with immune-mediated diseases of the kidney and the intestine. Our results identify ISR/eIF2α-mediated control of cytokine mRNA translation as an underlying mechanism that restricts Trm cell activity in homeostasis but also promotes rapid response upon local infection or autoimmune reaction.

## INTRODUCTION

T-cell memory is a hallmark of the adaptive immune system that results from the antigen-specific activation, expansion, and maintenance of T cells. Antigen-experienced memory T cells elicit a faster and more potent immune response upon re-exposure to the same antigen, providing superior protection against reinfection (*1*). Memory T cells were divided into two distinct populations, central memory T (Tcm) and effector memory (Tem) cells, which can be distinguished based on specific homing capacity and effector functions (*1*). In the last decade, tissue-resident memory T (Trm) cells were described as a distinct memory T cell subset (*2–6*). In contrast to Tcm and Tem cells, which survey peripheral tissues by recirculating through the blood and lymph, Trm cells do not recirculate and exhibit long-term persistence at sites of previous infections. It is now well established that Trm cells provide long-term immunity against local reinfections (*2–6*), but they also contribute to organ-specific immunopathology in relapsing immune-mediated inflammatory diseases of the gut (*7*), central nervous system (*8*), skin (*9, 10*), and kidney (*11–13*). The local production of cytokines by CD4^+^ Trm cells orchestrates protective tissue immunity against invading pathogens, but can also promote hyperinflammation and tissue damage in immune-mediated inflammatory diseases (*14–18*). A better understanding of the mechanisms that enable Trm cells to mount rapid local cytokine responses upon activation while restricting their cytokine production in steady-state could pave new ways for targeted treatment strategies for organ-specific autoimmunity and infection.

Here we show that Trm cells in healthy human organs (kidney, lung, and gut) contain mRNA coding for proinflammatory type-1 and type-3 cytokines in the absence of a detectable inflammatory tissue response. Translation of cytokine mRNA in CD4^+^ Trm cells is inhibited by activation of the integrated stress response (ISR)/eukaryotic translation initiation factor 2 alpha (eIF2α) pathway (*19*). Upon activation of Trm cells, eIF2α is dephosphorylated, leading to the immediate translation of the preformed cytokine mRNA. These findings suggest that the ISR/eIF2α pathway tightly controls effector cytokine production in Trm cells, identifying a hitherto unknown mechanism of Trm-cell regulation under homeostatic and inflammatory conditions.

## RESULTS

### CD4^+^ Trm cells express proinflammatory cytokine mRNA in healthy human tissues

Recent studies have revealed a significant number of memory T cells in non-lymphoid tissues under homeostatic conditions (*20*). Yet, limited information regarding the mechanisms controlling their function is available. To fill this gap, we comprehensively analyzed CD4^+^ and CD8^+^ T cells isolated from healthy renal tissue and matched blood samples (Figure 1A) (*13*). Using combined single-cell RNA sequencing (scRNA-seq) and cellular indexing of transcriptomes and epitopes by sequencing (CITE-seq), we examined the transcriptomic profile of T cells. We found that type-1 and type-3 proinflammatory cytokine mRNAs were expressed in T cells in the kidney under steady-state conditions. Corresponding T cells from the peripheral blood, by contrast, displayed much lower levels of these cytokine mRNAs (Figures 1B and C, and Supplementary Figures 1A and B). RNA fluorescence *in-situ* hybridization (FISH) analysis confirmed the presence of CD3^+^ T cells expressing *IL17A*, *IFNG,* and *CSF2* mRNA in healthy kidney tissue specimens (Figure 1D). Of note, the mRNA expression of the proinflammatory type-3 cytokines *IL17A* and *IL17F*, and to a lesser degree of *CSF2* and *TNF,* was higher in CD4^+^ T cells compared with CD8^+^ T cells in the kidney (Figure 1C). We, therefore, focused on renal CD4^+^ T cells in our further studies.

**Figure 1:**
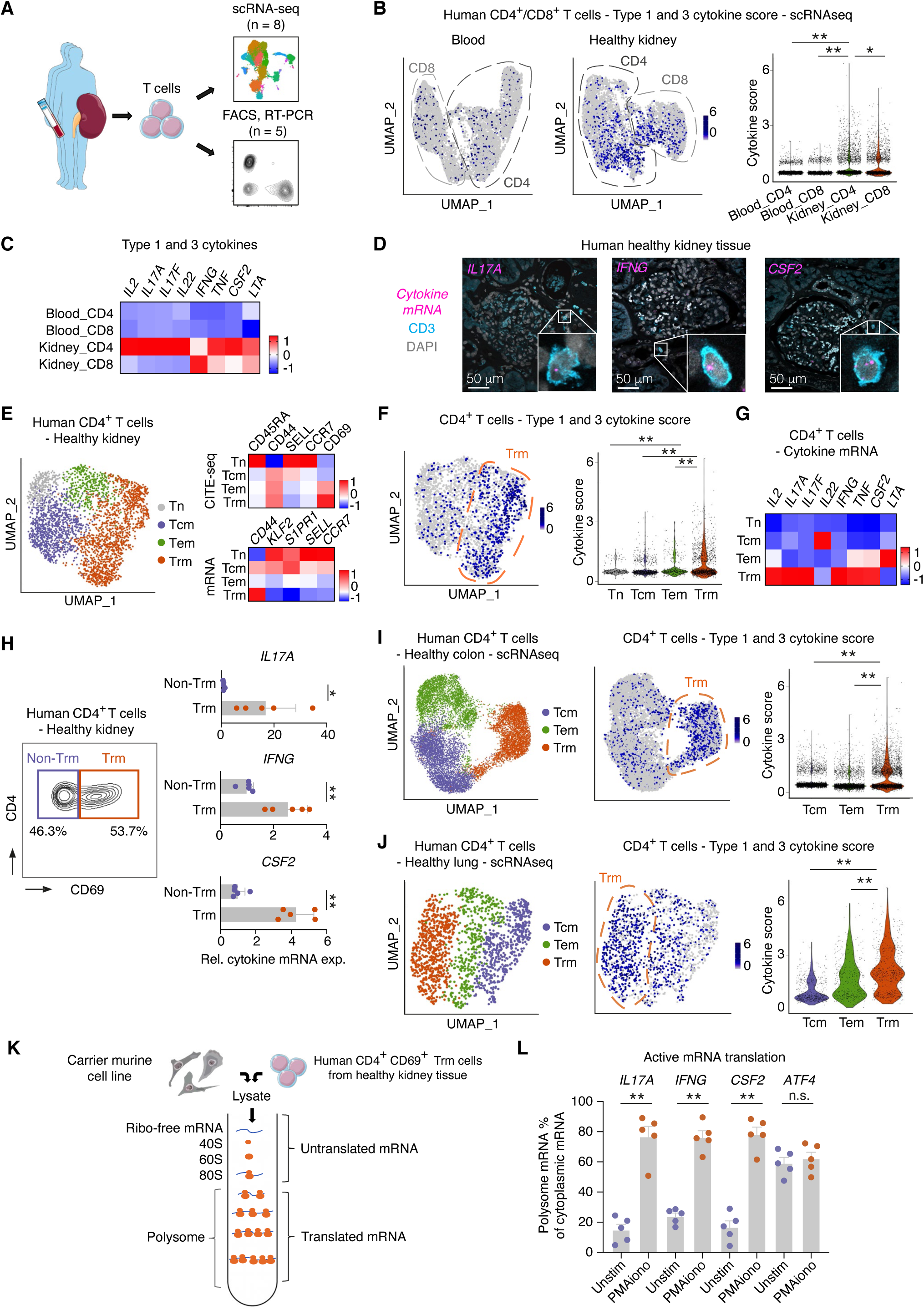
CD4^+^ Trm cells express proinflammatory cytokine mRNA without translation under homeostatic conditions. (A) Schematic overview of the experiment. FACS-sorted T cells isolated from healthy human kidney tissue and matched blood were analyzed using scRNA-seq (n = 8) and other analyses, including quantitative RT-PCR and sucrose gradient fractionation (n = 5). (B) UMAP plots of CD4^+^ and CD8^+^ T cells from peripheral blood and healthy kidney tissue displaying type-1 and type-3 cytokine scores. This score was calculated based on the expression of the genes listed in a heatmap depicted in C (C) Heatmap showing the expression of cytokine mRNA in CD4^+^ and CD8^+^ T cells isolated from the blood and healthy kidney tissue. (D) Representative images of cytokine mRNA FISH combined with immunofluorescence staining of CD3 in healthy human kidney tissue. (E) UMAP plot of CD4^+^ T cells from healthy human kidney tissue. Clusters were annotated according to their gene and protein expression profiles. (F) UMAP plot displaying the type-1 and type-3 cytokine scores. A violin plot of the cytokine score of each T-cell subset is also shown. (G) Heatmap showing type-1 and type-3 cytokine mRNA expression in different T-cell subsets. (H) Representative FACS plot and the result of RT-PCR analysis for the expression of *IL17A*, *IFNG*, and *CSF2* mRNA in CD4^+^ CD69^+^ Trm cells and CD4^+^ CD69^-^ non-Trm cells sorted from healthy human kidney tissue (n = 5). (I and J) UMAP plots of CD4^+^ T cells isolated from the healthy human colon (n = 4) (I) and lung (n = 10) (J). Clusters were annotated according to their gene expression profiles. UMAP plots and violin plots showing the type-1 and type-3 cytokine scores. (K) Schematic representation of the sucrose gradient centrifugation. Lysates of human renal CD4^+^ CD69^+^ Trm cells were subjected to centrifugation and separated into translated mRNA and untranslated mRNA fractions using the polysome profiling technique. (L) Cytokine and non-cytokine transcripts in translated (polysome) or untranslated (non-polysome) mRNA fractions were quantified by RT-PCR to calculate mRNA translation efficiency. Data are mean + S.E.M (* P < 0.05, ** P < 0.01).

Unsupervised clustering of CD4^+^ T cells from the human kidney identified naïve T (Tn), Tcm, Tem, and Trm cells (Figure 1E and Supplementary Figure 1C). CD4^+^ Trm cells represented the predominant cellular source of proinflammatory cytokines (Figures 1F and G and Supplementary Figure 1C). This finding was further corroborated by quantitative RT-PCR analysis, which demonstrated that proinflammatory cytokine mRNA level was significantly higher in CD4^+^ CD69^+^ Trm cells compared with CD4^+^ CD69^-^ non-Trm cells isolated from healthy renal tissue (Figure 1H).

To explore whether homeostatic cytokine mRNA expression in CD4^+^ Trm cells is specific to the kidney or an overarching mechanism shared between different organs, we analyzed published scRNA-seq data from the colon (*21*) and lung (*22*) of healthy individuals. Consistent with the findings in the kidney, CD4^+^ Trm cells were present in the healthy colon and lung, and these cells expressed the highest levels of proinflammatory cytokine mRNA compared with other T-cell populations (Figures 1I and J, and Supplementary Figure 2). These data show that Trm cells express proinflammatory cytokines in various human tissues under non-inflammatory conditions.

### Cytokine mRNA in CD4^+^ Trm cells is not translated under homeostatic conditions

Uncontrolled cytokine production might lead to inflammation and immunopathology. In healthy human kidney tissue, we did not observe signs of active inflammation despite the presence of cytokine mRNA-expressing Trm cells (Supplementary Figure 3A). Therefore, we sought to investigate whether proinflammatory cytokine mRNA expression in T cells gives rise to tissue responses under homeostatic conditions. To that end, we first analyzed the European Renal cDNA Bank database (*23*) to examine cytokine-responsive gene expression in health and disease. This dataset encompassed biopsies derived from kidneys of healthy individuals (living donor kidney transplantation) and from patients with autoimmune kidney diseases such as anti-neutrophil cytoplasmic antibody-associated glomerulonephritis (ANCA-GN), lupus nephritis, and IgA nephropathy. Notably, nephritic tissue exhibited increased cytokine-responsive gene expression and cytokine tissue scores, as determined by genes induced by IL-17, INF-γ, or GM-CSF (*24–26*) (Supplementary Figures 3B and C). In contrast, the expression of cytokine-responsive genes was significantly lower in healthy kidney tissue (Supplementary Figures 3B and C) irrespective of the homeostatic cytokine mRNA expression by T cells (Figures 1A-G). These results suggest that cytokine mRNA in CD4^+^ Trm cells does not lead to tissue responses under homeostatic conditions.

To gain further insight into the translational regulation of cytokine mRNAs, we employed the sucrose gradient centrifugation and polysome profiling technique (*27*) which allowed us to separate actively translated mRNAs from those that were not translated. We isolated CD4^+^ CD69^+^ Trm cells from healthy human kidney tissue, lysed the cells to preserve intact mRNA-ribosome complexes and, following sucrose gradient centrifugation, separated translated (i.e. polysome) and untranslated mRNA to quantify them using RT-PCR analysis (Figure 1K). When Trm cells were not stimulated, most *IL17A*, *IFNG*, and *CSF2* mRNAs were detected in the untranslated fraction, indicating that the translation of proinflammatory cytokine mRNA is suppressed under homeostasis (Figure 1L). However, after stimulation of Trm cells with PMA/ionomycin for one hour, most of the cytokine mRNA was detected in the polysome fraction of the translated mRNAs (Figure 1L). In contrast, *ATF4* mRNA, which is translated under compromised cap-dependent translation (*19*), was efficiently translated without prior T-cell stimulation (Figure 1L), indicating that the translation of cytokine mRNAs is actively suppressed in CD4^+^ Trm cells under homeostatic conditions.

### mRNA translation pathway is suppressed in CD4^+^ Trm cells

To study the regulation of mRNA translation in Trm cells, we took advantage of a murine model system. Because laboratory mice maintained under specific pathogen-free (SPF) conditions are exposed to a limited number of microbes and pathogens, they harbor only a few memory T cells in non-barrier organs such as the kidney. Therefore, we adopted an experimental Trm-cell induction model (*13, 28*) for our studies (Figure 2A). In this model, mice were systemically infected with *Staphylococcus aureus* and then treated with antibiotics to clear the infection which efficiently induced renal IL-17A-producing Trm cells (Trm17 cells) (*13*). Immunofluorescence staining combined with mRNA FISH at two months after clearance of the infection, a time point at which no tissue inflammation or pathology is detectable, revealed the presence of numerous T cells in the kidney (Figure 2B), some of which expressed proinflammatory cytokine mRNA.

**Figure 2:**
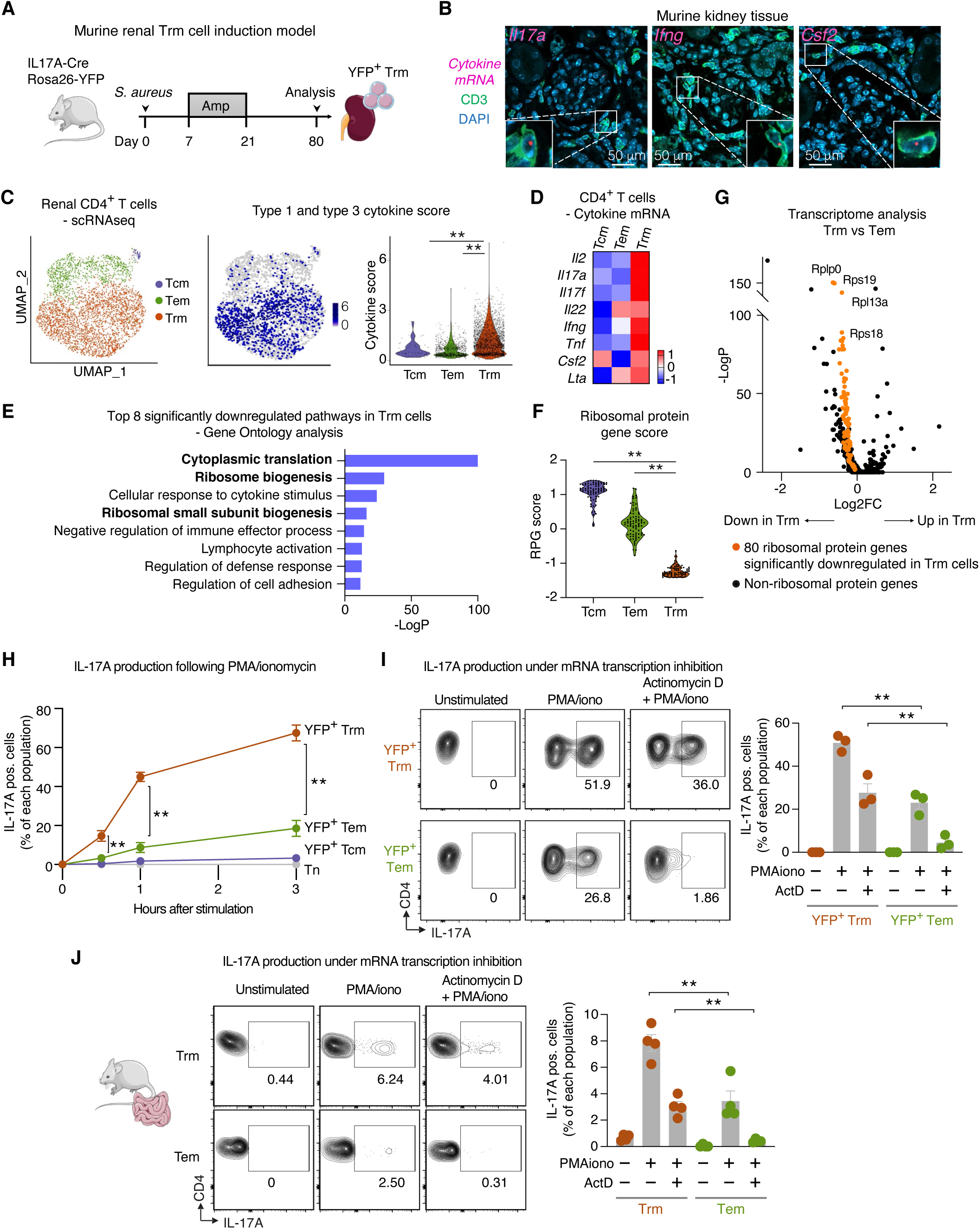
Murine Trm cells show downregulated mRNA translation profile under homeostasis with immediate cytokine production ability upon stimulation. (A) Schematic representation of the murine Trm-cell induction model. Wildtype or IL-17A fate reporter mice were infected with *S. aureus*, treated with ampicillin, and analyzed at least 2 months after initiating the administration of antibiotics. (B) Representative images of cytokine mRNA FISH combined with immunofluorescence CD3 staining of kidney sections from mice with prior Trm-cell induction. (C) scRNA-seq of CD4^+^ YFP^+^ T cells sorted from kidneys of the Trm-cell induction model using IL-17A fate reporter mice. Clusters were annotated according to their gene expression profiles. The type-1 and type-3 cytokine scores were calculated by the expression of genes listed in a heatmap in D. Violin plot of the cytokine score of each T-cell population is also shown. (D) Heatmap showing proinflammatory cytokine gene expression in different T-cell subsets. (E) Gene Ontology analysis displaying pathways downregulated in Trm cells compared with Tem cells. (F) Violin plot showing the ribosomal protein gene score (based on the genes in the heatmap in Supplementary Figure 4D) for each murine T-cell subset. (G) Volcano plot showing differential gene expression between murine Trm and Tem cell clusters. Ribosomal protein genes downregulated in Trm cells are shown in orange. (H) IL-17A production by murine renal T cells. T cells were stimulated for 30 min, 1 hour, and 3 hours with PMA/ionomycin (n=6). (I) IL-17A production by T cells treated with PMA/ionomycin with or without 20 ug/mL actinomycin D (transcription inhibitor) for 4 hours (n = 3). (J) CD4^+^ Trm and Tem cells from the small intestine of wildtype mice were stimulated by PMA/ionomycin with or without 20 ug/mL actinomycin D, and IL-17A production was analyzed (n = 4). Data are mean + S.E.M (** P < 0.01).

To characterize the transcriptome of murine renal T cells, we sorted CD4^+^ YFP^+^ T cells from the kidneys using IL-17A fate reporter mice (*Il17a*^Cre^ × *R26*^eYFP^) and performed scRNA-seq analysis. CD4^+^ T cell clustering based on gene expression profile identified Tcm, Tem, and Trm cells (Figure 2C and Supplementary Figure 4A). As seen in human CD4^+^ T cells, murine Trm cells demonstrated expression of proinflammatory cytokine mRNA (Figures 2C and D, and Supplementary Figure 4B). The *Il17a* mRNA expression in renal Trm17 cells was further confirmed by RT-PCR (Supplementary Figure 4C).

To unravel the molecular pathways active in Trm cells under non-inflammatory conditions, we conducted the Gene Ontology pathway analysis using the scRNA-seq data. Pathways associated with cytoplasmic translation and ribosome biogenesis were significantly downregulated in Trm cells compared with Tem cells (Figure 2E). In line with this result, we observed coordinated downregulation of genes coding for ribosomal proteins in Trm cells (Figures 2F and G, and Supplementary Figure 4D), indicating suppression of the global mRNA translation system in Trm cells. Moreover, the protein-protein interaction enrichment analysis (*29*) exhibited the suppression of the ribosomal scanning and start codon recognition processes – crucial steps in translation initiation (Supplementary Figure 4E). These findings demonstrate a significant suppression of mRNA translation in Trm cells while expressing proinflammatory cytokine mRNA under homeostatic conditions.

### Already transcribed cytokine mRNA is immediately available for translation in murine Trm cells

The rapid production of cytokines is crucial to mount an effective host defense. We aimed to investigate the kinetics of cytokine production upon reactivation of memory T-cell populations. To this end, T cells were isolated from the kidneys of the experimental Trm-cell induction model and stimulated these cells with PMA/ionomycin. Cytokine production was assessed at different time points. We found that Trm cells produced IL-17A and other cytokines faster than Tcm-cell and Tem-cell subsets (Figure 2H and Supplementary Figure 4F).

To address whether the already transcribed cytokine mRNA fosters a more rapid and stronger cytokine production by Trm cells, we isolated the Trm17 and Tem17 subsets from the kidneys of IL-17A fate reporter mice. We then stimulated them with PMA/ionomycin with or without actinomycin D, which blocks mRNA transcription. Trm cells and, to a lesser extent, Tem cells produced IL-17A protein upon stimulation without actinomycin D (Figure 2I), corresponding with higher *Il17a* mRNA expression levels in Trm cells (Figures 2C and D). When Trm cells were stimulated in the presence of actinomycin D, they still produced IL-17A protein (Figure 2I), indicating that already transcribed *Il17a* mRNA pools were readily available for translation. In contrast, IL-17A protein production by Tem cells was almost completely abolished by actinomycin D treatment, as this T-cell subset contains low mRNA levels (Figures 2C and D). To verify our findings, we also examined CD4^+^ Trm and Tem cells isolated from the small intestine of healthy wildtype mice and obtained identical results (Figure 2J). These findings demonstrate that cytokine mRNA depots in Trm cells are sufficient for immediate cytokine protein production upon reactivation, explaining the rapid and pronounced cytokine response in this T-cell population.

### ISR/eIF2α pathway is activated in human Trm cells under homeostatic conditions

To understand the molecular pathways regulating mRNA translation in Trm cells, we performed a comprehensive analysis of scRNA-seq data from different human organs, by using a combination of data generated for this study (see Fig. 1) and published data sets (*21, 22*). Similar to the mouse model, the GO term pathway analysis revealed downregulation of the translation and ribosome biogenesis pathways in Trm cells obtained from the healthy human kidney (Fig. 3A and B) and other healthy tissues (colon and lung, Supplementary Figures 5A-E). In addition, the Reactome pathway analysis revealed an upregulation of the cellular responses to stress pathway in these cells (Figure 3C). This pathway is associated with the ISR, which is a biological process activated by various stimuli such as hypoxia, heat stress, cell starvation, and unfolded protein response (*19*). The ISR downregulates global mRNA translation through the phosphorylation of eIF2α, which markedly decreases the cap-dependent translation initiation (*19*). Indeed, the downregulation of the start codon recognition and translation initiation process was also observed in murine scRNA-seq data analysis (Supplementary Figure 4E). Consistent with these findings, human Trm cells exhibited the highest expression of ISR-associated genes (*19, 30–34*) (Figure 3D and Supplementary Figures 5F-I). We observed phosphorylation of eIF2α under unstimulated conditions and immediate dephosphorylation eIF2α upon Trm cell stimulation (Figure 3E and Supplementary Figures 5J and K). In line with this, a puromycin uptake assay revealed higher levels of the global protein synthesis in stimulated Trm cells (Supplementary Figure 5L).

**Figure 3:**
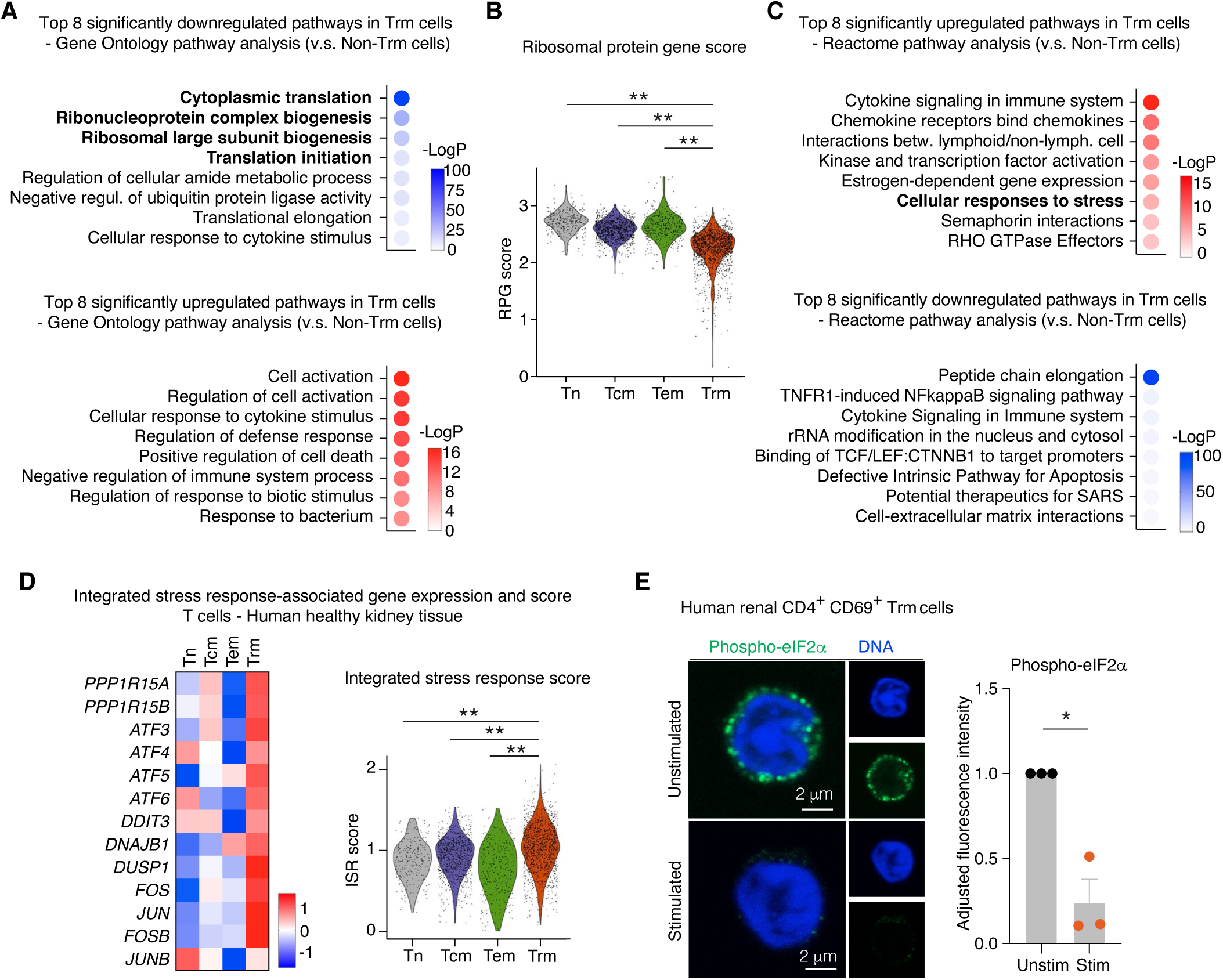
The integrated stress response pathway is activated in human Trm cells. (A) Gene Ontology analysis showing pathways downregulated in Trm cells compared with non-Trm cells from human healthy kidney tissue. (B) Ribosomal protein gene score (based on the same genes in Figure 2F) in each T-cell subset. (C) Reactome analysis displaying pathways upregulated in Trm cells compared with non-Trm cells. (D) Expression of the integrated stress response (ISR)-associated genes displayed as a heatmap, and ISR-score calculated based on the genes shown in the heatmap. (E) Representative images of phospho- and total eIF2α staining in sorted Trm cells with or without PMA/ionomycin stimulation. Quantification for the fluorescence intensity is also shown. Data are mean + S.E.M (* P < 0.05, ** P < 0.01).

### Cytokine production is controlled by the ISR/eIF2α pathway in murine Trm cells

To investigate the role of ISR in T-cell function, we utilized murine Trm cells. Similar to human Trm cells, unstimulated murine renal Trm cells displayed high levels of eIF2α phosphorylation (Figure 4A). Upon stimulation with PMA/ionomycin or anti-CD3/28 Ab, eIF2α was dephosphorylated, leading to increased global protein synthesis (Figure 4A and Supplementary Figures 6A - C). Murine Trm cells showed higher phospho-eIF2α levels than Tem and Tcm cells (Supplementary Figure 6D). Following ISR activation, untranslated mRNAs are sequestered into stress granules (SGs), which are membrane-less cytoplasmic organelles (*27*). mRNA FISH analysis for IL-17A combined with immunocytochemistry for the SG marker Tia1 revealed colocalization of cytoplasmic *Il17a* mRNA with the SG marker Tia1 in unstimulated Trm cells (Figure 4B). In contrast, Trm-cell stimulation reduced the association of *Il17a* mRNA with SGs (Figure 4B), indicating the release of mRNA for translation.

**Figure 4:**
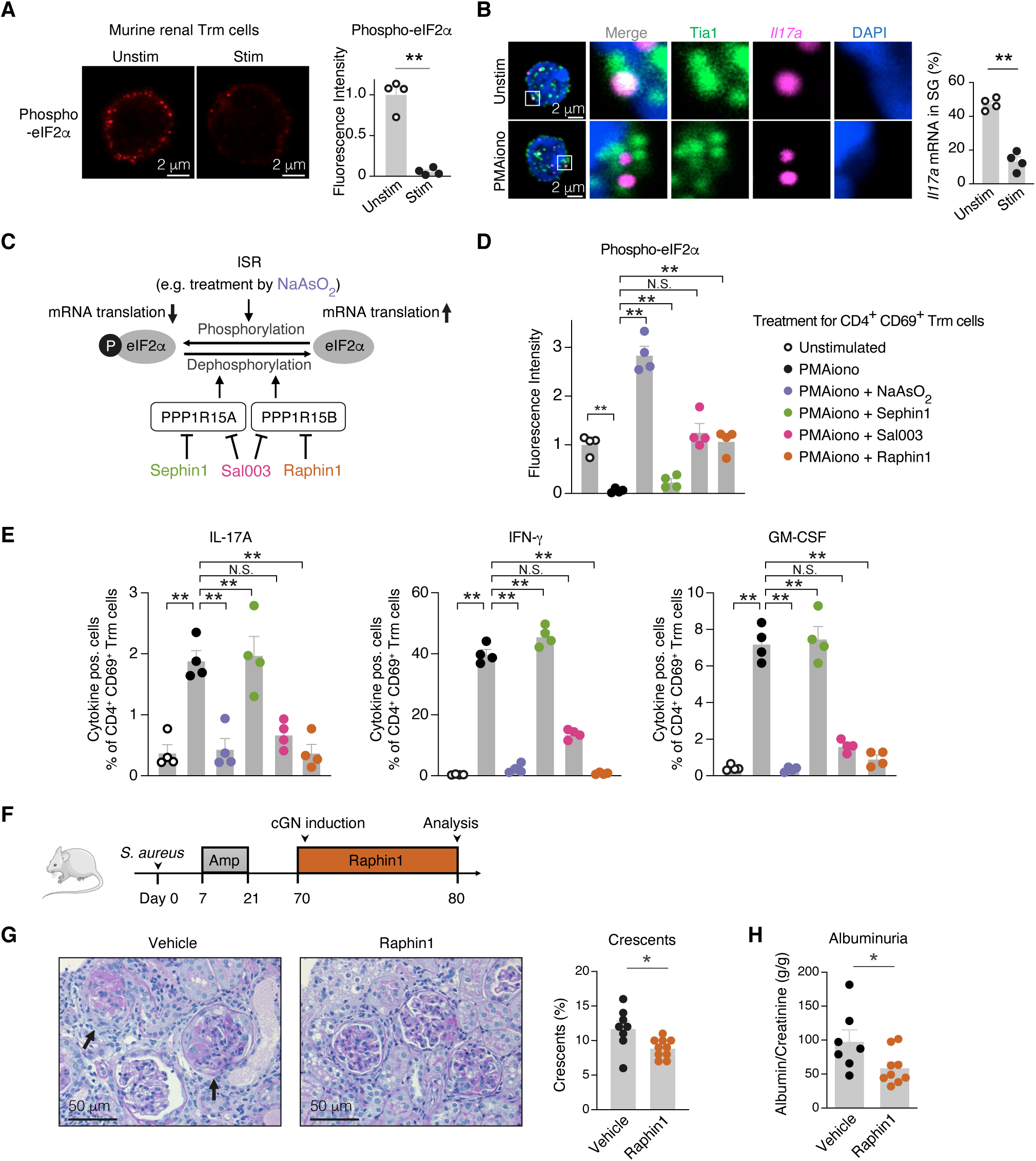
The integrated stress response pathway regulates cytokine production. (A) Representative images of phospho- and total eIF2α immunocytochemistry in murine renal Trm cells before and after stimulation with PMA/ionomycin. Quantification is also shown (n = 4). (B) mRNA FISH for *Il17a* mRNA and immunocytochemistry for Tia1 (stress granule marker) before and after stimulation with PMA/ionomycin. Representative images and a quantification result (n = 4) are shown. (C) Schematic representation of eIF2α regulation. eIF2α is phosphorylated upon activation of the integrated stress response pathway (e.g., by treatment with NaAsO_2_). Phosphatases for eIF2α can be blocked by the non-selective inhibitor Sal003, or by the selective inhibitors Sephin1 or Raphin1 targeting phosphatases PPP1R15A or PPP1R15B, respectively. (D and E) Phospho-eIF2α levels (D) and frequency of cytokine^+^ cells (E) in total Trm cells stimulated by PMA/ionomycin in the presence of indicated compounds (n = 4). Trm cells were treated with vehicle, 500 μM NaAsO_2_, 10 μM Sal003, 20 μM Sephin1, or 20 μM Raphin1 for 3 hours. (F) Schematic representation of the experimental glomerulonephritis model. Mice were infected with *S. aureus* and treated with ampicillin to induce kidney Trm cells. Two months later, crescentic glomerulonephritis (cGN) was induced, and an eIF2α phosphatase inhibitor, Raphin1, was given twice daily by oral gavage between d70 and d80. Kidney function was evaluated 10 days after cGN induction. (G) Representative photographs of PAS-stained kidney sections and quantification of glomerular crescents from vehicle and Raphin1-treated mice. Arrows indicate glomeruli with glomerular crescents and necrosis. (H) Quantification of albuminuria. The *in vivo* data are representative of two independent experiments. Data are mean + S.E.M (* P < 0.05, ** P < 0.01).

### eIF2α phosphorylation constrains cytokine mRNA translation in Trm cells

To further investigate the role of eIF2α phosphorylation in inhibition of cytokine production, we treated Trm cells with sodium arsenite, commonly used to mimic ISR activation by oxidative cues (*27*) (Figure 4C). Sodium arsenite treatment resulted in marked eIF2α phosphorylation (Figure 4D) and suppressed the production of IL-17A, IFN-γ, and GM-CSF from activated Trm cells (Figure 4E) without causing any noticeable cell toxicity (Supplementary Figure 6E). Together, our results suggest that ISR and eIF2α mediate the translation status of the proinflammatory cytokines. Phosphorylated eIF2α and activated ISR suppress translation by sequestration of the proinflammatory cytokines mRNA in stress granules and vice versa, dephosphorylation of eIF2α and ISR suppression activates their translation.

Since eIF2α was phosphorylated in Trm cells under steady state conditions, we hypothesized that blocking eIF2α dephosphorylation might be sufficient to maintain eIF2α phosphorylation levels and inhibit cytokine mRNA translation in activated Trm cells. As shown in Figure 4C, two phosphatase complexes (PPP1R15A-PP1 and PPP1R15B-PP1) dephosphorylates eIF2α and counteract ISR-induced inhibition of protein synthesis (*19*). The pan-eIF2α phosphatase inhibitor Sal003 (Figure 4C) significantly suppressed eIF2α dephosphorylation and consequently, the cytokine production in activated Trm cells (Figures 4D and E, and Supplementary Figure 6F), supporting the notion that eIF2α dephosphorylation is a necessary step for cytokine production in activated Trm cells. Since Sal003 blocks both eIF2α phosphatases, we tested other selective inhibitors Sephin1 and Raphin1, which block PPP1R15A and PPP1R15B, respectively (Figure 4C). Sephin1 did not suppress eIF2α dephosphorylation or cytokine protein production. In contrast, Raphin1 significantly reduced eIF2α dephosphorylation and potently suppressed cytokine production in Trm cells (Figures 4D and E, and Supplementary Figure 6F). These findings demonstrate that eIF2α dephosphorylation is a crucial step in the initiation of rapid translation of cytokine mRNAs in Trm cells and is predominantly driven by PPP1R15B.

### Targeting eIF2α dephosphorylation ameliorates experimental autoimmune glomerulonephritis

To investigate the potential therapeutic effects of targeting eIF2α dephosphorylation in autoimmune disease, we used a well-established preclinical murine model of autoimmune glomerulonephritis (crescentic GN) (*13*). After induction of renal Trm cells, we challenged mice with crescentic GN (cGN) and treated these animals with the orally available eIF2α phosphatase PPP1R15B inhibitor Raphin1 (*35*) (Figure 4F). As previously reported (*35*), Raphin1 was bioactive and augmented the levels of phospho-eIF2α in Trm cells *in vivo* (Supplementary Figure 7A). Most importantly, Raphin1 treatment ameliorated cGN course regarding glomerular crescent formation and albuminuria compared to the control group (Figures 4G and H). These findings suggest that targeting the ISR/eIF2α pathway may hold promise as a therapeutic strategy for treating immune-mediated diseases.

### The ISR pathway is downregulated in patients with immune-mediated glomerular diseases and inflammatory bowel disease

Finally, we investigated the relevance of our findings in human autoimmune kidney disease by analyzing scRNA-seq data of renal CD4^+^ T cells obtained from patients with ANCA-GN and healthy controls (Figure 5A and B). The mRNA expression levels of type-1 and type-3 cytokines in Trm cells from ANCA-GN patients were similar to those in healthy controls (Figure 5C). We conducted the Reactome pathway analysis comparing Trm cells from ANCA-GN patients and controls. Of note, these analyses indicated a downregulation of the cellular response to stress pathway in Trm cells from ANCA-GN patients (Figure 5D). Moreover, Trm cells from the ANCA-GN group exhibited a significantly lower ISR-associated gene score (Figure 5E and Supplementary Figure 8A). These findings would indicate a higher mRNA translation efficiency in Trm cells from ANCA-GN patients. This finding is noteworthy because cytokine-responsive genes were highly expressed in kidney tissue from patients with ANCA-GN compared with healthy controls (Figure 5F), suggesting active cytokine mRNA translation as a mechanism of T-cell mediated tissue damage in glomerulonephritis (*13, 36*). Furthermore, we extended our analysis to include scRNA-seq data obtained from colon CD4^+^ T cells derived from healthy controls and patients with Crohn’s disease, an inflammatory bowel disease in which type-1 and type-3 cytokines play key roles (*37*). In this published dataset, we identified two Trm cell clusters, namely the *TNF*-high Trm cluster and the *IL17A*-high Trm cluster (Figure 5G). As observed in the ANCA-GN dataset, the expression levels of inflammatory cytokine mRNA in Trm cells were comparable between the healthy controls and Crohn’s disease patients (Figure 5H). Both Trm cell populations exhibited a downregulation of the cellular response to stress pathway and a lower ISR score in patients with Crohn’s disease as compared to the healthy control (Figure 5I and J, Supplementary Figure 8B). These findings underscore the potential role of downregulating the ISR pathway as a mechanism that enhances the synthesis of effector molecules, resulting in increased tissue inflammation.

**Figure 5:**
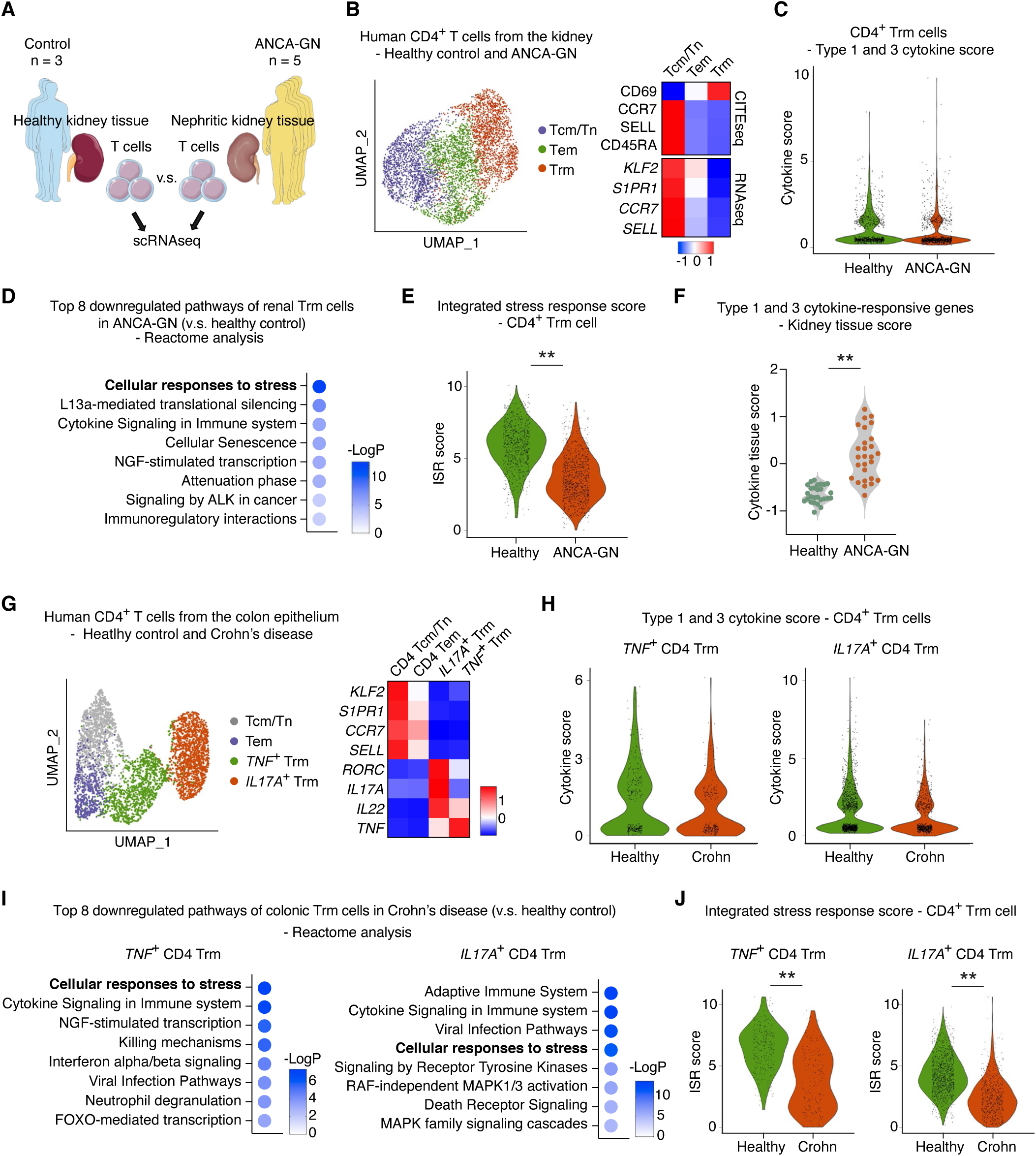
The integrated stress response pathway is downregulated in Trm cells in patients with inflammatory diseases. (A) T cells isolated from kidney biopsies of patients with ANCA-GN were compared with those from healthy kidney tissue using scRNA-seq to investigate differences in gene expression profiles. (B) UMAP plot of renal CD4^+^ T cells from healthy controls and ANCA-GN patients. Clusters were annotated according to their gene and protein expression profiles. (C) The type-1 and type-3 cytokine scores (based on the same genes as Figure 1B) in renal Trm cells from healthy control and ANCA-GN patients. (D) Reactome analysis showing downregulated pathways in CD4^+^ Trm cells from ANCA-GN patients compared with the control group. (E) ISR score (based on the same genes as Figure 3D) of CD4^+^ Trm cells from ANCA-GN patients and controls. (F) Type-1 and type-3 cytokine-responsive gene score was calculated using European Renal cDNA Bank transcriptome data generated from kidney biopsies (based on supplementary Figures 3B and C). (G) UMAP plot of colon CD4^+^ T cells from healthy controls and patients with Crohn’s disease. Clusters were annotated according to their gene expression profiles. (H) The type-1 and type-3 cytokine scores in colon Trm cells from healthy control and Crohn’s disease patients. (I) Reactome analysis showing downregulated pathways in CD4^+^ Trm cells from Crohn’s disease patients compared with the control group. (J) ISR score (based on the same genes as Figure 3D) of CD4^+^ Trm cells from Crohn’s disease patients and control. Data are mean + S.E.M (* P < 0.05, ** P < 0.01).

## DISCUSSION

Trm cells provide rapid protection against local infections. Yet, they also promote tissue damage in autoimmunity. Therefore, a deeper insight into the mechanisms controlling Trm-cell function is critical. Here, we demonstrate that CD4^+^ Trm cells embody a poised cell population, which stores translationally suppressed proinflammatory type-1 and type-3 cytokine mRNA under homeostatic conditions. This enables the immediate translation and production of proinflammatory cytokines upon Trm-cell activation. We further identify the ISR and its mediator eIF2α as a central node in facilitating the storage of cytokine mRNA in stress granules and thereby suppressing its translation.

mRNA expression profiling of T cells in healthy human organs, such as the kidney, gut, and lung, revealed that CD4^+^ Trm cells express mRNAs for proinflammatory cytokines. Expression of cytokine mRNAs without an apparent inflammatory response suggests that the mRNAs are not translated to cytokines under homeostatic conditions. The presence of preformed cytokine mRNA in T cells under homeostatic conditions was noted in previous studies (*38–43*), however, the mechanism of translational repression and its relevance for tissue-specific immunity in health and disease remained elusive. Our results reveal that pathways associated with global mRNA translation, ribosomal biogenesis, and mRNA translation initiation are significantly suppressed in CD4^+^ Trm cells in homeostasis, suggesting that attenuation of mRNA translation is major mechanism to restrict Trm cell activity and thereby prevent tissue inflammation. Our results further link the control of translation in Trm cells to the ISR pathway, an evolutionarily conserved mechanism allowing cells to adapt to different environments by reprogramming gene expression to reduce global mRNA translation and ribosome biogenesis (*19, 44*). Various physiological or pathological stress stimuli including hypoxia, amino acid deprivation, glucose/nutrition deprivation and viral infection are sensed by four specialized kinases (PERK, GCN2, PKR and HRI) that each phosphorylate Ser^51^ in eIF2α to activate the ISR (*45*). Two specific phosphatases counteract this reaction by promoting the dephosphorylation of eIF2α. Both phosphatases contain a common core catalytic subunit, protein phosphatase 1, and a specific regulatory subunit (PPP1R15A or PPP1R15B) that confers the phosphatase specificity to eIF2α (*46, 47*). In contrast to the emerging roles of the ISR pathway in brain health and cognitive disorders (*19*) (e.g. inhibition of the ISR enhances long-term memory formation), little is known about the function of this pathway in the immune system.

A potential role of the ISR/eIF2α axis in T-cell biology was first suggested over a decade ago based on *in vitro*-polarized T cells in which eIF2α was phosphorylated under steady-state conditions. Stimulation of these T cells resulted in rapid eIF2α dephosphorylation, ribosomal mRNA loading, and cytokine secretion (*43*). The role of ISR in the maintenance of naïve T-cell homeostasis was also reported (*48*). Here, we show that Trm cells in non-inflamed peripheral tissues maintain an activated ISR/eIF2α pathway and provide evidence that augmented ISR inhibits the translation of cytokine mRNA stored in stress granules. Stimulation of Trm cells via the T cell receptor signaling leads to the dephosphorylation of eIF2α mainly by the phosphatase PPP1R15B and a rapid activation of translation of the proinflammatory cytokine mRNAs. Pharmacological PPP1R15B inhibition in an experimental mouse model of autoimmune kidney disease resulted in less severe glomerulonephritis in terms of glomerular crescent formation and proteinuria. Furthermore, we demonstrate that the ISR pathway is downregulated in human Trm cells in autoimmune kidney disease and inflammatory bowel disease, in which cytokine production, e.g. of IL-17A, IFN-γ and GM-CSF, mediates tissue damage (*13, 36, 37*) (Figures 5A-J), thus further suggesting the importance of this pathway in inflammatory diseases.

Cytokine mRNA translation needs to be controlled to prevent excessive cytokine production. To date, different mechanisms of translation regulation have been reported. How the ISR pathway synergizes with other mRNA translation regulation mechanisms, e.g., RNA-binding proteins (*42, 49*) or the mTOR pathway (*50*), remains elusive. The precise mechanisms and kinases that activate the ISR pathway in Trm cells in tissue homeostasis and how an inflammatory environment reverses the ISR/eIF2α pathway activation in Trm cells remain to be fully elucidated. It is plausible, however, that their extravascular location in hypoxic areas of the kidney (*51, 52*) and other organs (*53, 54*) could serve as one of the triggers for ISR in Trm cells, given that hypoxia is one of the stimuli for eIF2α phosphorylation (*55*). Of note, canonical pathway analysis of the scRNA-seq data of T cells from human healthy kidney revealed that hypoxia-associated pathways were upregulated in Trm cells (Supplementary Figure 9).

Overall, our findings suggest that the ISR/eIF2α pathway acts as a mechanism regulating the activation and cytokine secretion of Trm cells in homeostasis and inflammation, and identify this pathway as a potential therapeutic target in T-cell-mediated inflammatory diseases.

## METHODS

### Human studies

Human kidney tissue was obtained from patients included in the Hamburg GN Registry, the European Renal cDNA Bank, and the Clinical Research Unit 228 (CRU 228) ANCA-GN cohort. In some instances, matched blood samples from the same patients were also analyzed for this study. The healthy part of the kidney, which was removed due to tumor nephrectomy, was dissected to analyze renal T cells under homeostatic conditions. The histological examination confirmed that the healthy part of the kidney did not exhibit any signs of tumor invasion or active immune cell invasion. The research studies were conducted with the approval of the *Ethik-Kommission der Ärztekammer Hamburg* (the local ethics committee of the Chamber of Physicians in Hamburg) and in compliance with the ethical principles outlined in the Declaration of Helsinki. All participating patients gave informed consent for the research.

### Animal experiments

Experiments with mice followed the national guidelines, and local ethics committees approved the research protocols. Mice were housed under specific pathogen-free conditions to ensure the validity and reliability of the experiments. Male mice on a C57BL/6 background were used for all experiments and were age-matched to minimize the variability between subjects. Experimental crescentic glomerulonephritis was induced by i.p. administration of sheep serum targeted against the glomerular basement membrane, as previously described (*13*). *Il17a*^Cre^ × *R26*^eYFP^ mice (*56*) were used in this study to identify Trm17 cells. For the Raphin1 experiment, the compound was reconstituted in distilled water with 0.5% methylcellulose (Sigma, M0512) and administered twice a day by oral gavage. To induce *S. aureus* infection in mice, the *S. aureus* strain (SH1000) (*57*) was utilized. To initiate the infection, mice were administered 1×10^7^ staphylococci in 100 μl sterile PBS via intravenous injection.

### Histopathology, immunohistochemistry, and immunofluorescence

To evaluate glomerular crescent formation, 30 glomeruli per mouse were assessed in a blinded manner in periodic acid-Schiff (PAS)-stained paraffin sections of kidneys (*13*). For immunofluorescence staining, primary antibodies, including CD3 (A0452, Dako, Glostrup, Denmark), Phospho-eIF2α (MA5-32021, Invitrogen), total eIF2α (D7D3, Cell Signaling) and Tia1 (ab140595, Abcam) were incubated in blocking buffer at 4°C for 1 hour. Following PBS washing, fluorochrome-labeled secondary antibodies were applied, and the staining was visualized using an LSM800 with Airyscan and the ZenBlue software (all Carl Zeiss, Jena, Germany).

To detect mRNA (FISH) in mice and human kidney sections, RNAscope Multiplex Fluorescent Assay (Advanced Cell Diagnostics, Newark, CA) was employed. mRNA detection (FISH) in sorted T cells was carried out using the ViewRNA Cell Plus Assay kit (Cat. No. 88-19000, Invitrogen). The slides were imaged using a Zeiss LSM800 confocal microscope and analyzed with ZEN software (Carl Zeiss, Jena, Germany). To evaluate *Il17a* mRNA in stress granules, the percentage of *Il17a* signals overlapping with Tia1-positive granules was calculated.

### Isolation and flow cytometric analysis of human and murine leukocytes

Single-cell suspensions were obtained from kidney and blood samples to isolate and analyze human leukocytes. Kidney tissue was enzymatically digested with collagenase D at 0.4 mg/ml (Roche, Mannheim, Germany) and DNase I (10 μg/ml, Sigma-Aldrich, Saint Louis, MO) in RPMI 1640 medium at 37° C for 30 minutes, followed by dissociation with gentleMACS (Miltenyi Biotec). Blood samples were separated using Leucosep tubes (Greiner Bio-One, Kremsmünster, Austria). Samples were filtered through a 30-μm filter (Partec, Görlitz, Germany) prior to antibody staining and flow cytometry.

Cells from murine spleens were isolated by squashing the organ through a 70-µm cell strainer. Erythrocytes were lysed using a lysis buffer (155 mM NH_4_Cl, 10 mM KHCO_3_, 10 µM EDTA, pH 7.2). To isolate renal lymphocytes from mice, kidneys were enzymatically digested with 400 µg/ml collagenase D (Roche) and 10 U/ml DNase I (Sigma-Aldrich) at 37°C for 30 min. Subsequently, leukocytes were isolated by density gradient centrifugation using 37% Easycoll (Merck Millipore) and a filtration step using a 30-µm cell strainer (Partec). T cell isolation from the intestine is described previously. Briefly, murine intestine was cut longitudinally after removing Peyer’s patches and adventitial fat. To collect intraepithelial lymphocytes, the intestine tissue was incubated in HBSS containing 1mM dithioerythritol followed by a dissociation step using 1 mM EDTA for 20 min at 37° C. To collect lamina propria lymphocytes, the tissue was cut into small pieces and incubated for 45 min at 37° C in HBSS supplied with 1 mg/ml collagenase and 10 U/ml DNase I. Leukocytes were further enriched by Percoll gradient. Intraepithelial cells and lamina propria lymphocytes were pooled for analysis. Cells were surface stained with fluorochrome-conjugated antibodies (human: CD45 (H130), CD3 (OKT3), CD4 (RPA-T4), CD8 (RPA-T8), CD69 (FN50), CD45RA (HI100), GM-CSF (VD2-21C11), IL-17A (BL168), IL-17F (SHLR17), IFN-γ (4S.B3), TNF-α (MAB11), CCR7/CD197 (Go43H7); mouse: CD45 (30-F11), CD3 (145-2C11), CD4 (RM4-5), CD8 (53-6.7), GM-CSF (MP1-22E9), IL-17A (TC11-18H10), IL-17F (9D3.1C8), IL-22 (poly 5164), IFN-γ (XMG1.2), TNF-α (MP6-XT22); all BD Biosciences, Biolegend, or eBiosciences) and a fixable dead-cell stain (Molecular Probes) to exclude dead cells from the analysis. For intracellular staining, samples were processed using Cytofix/Cytoperm (BD Biosciences) according to the manufacturer’s instructions.

To assess the phospho-eIF2α and total eIF2α levels, cells underwent surface staining and were subsequently fixed and permeabilized using Foxp3/Transcription Factor Staining Buffer (eBioscience). The cells were then incubated with primary antibodies directed against phospho-eIF2α or total eIF2α (1:1000) for 40 minutes. After two rounds of washing, the cells were exposed to a secondary antibody conjugated with AF647 against Rabbit IgG (1:1000) for 40 minutes.

### Flow cytometry and cell sorting

Samples were measured with LSR II or Symphony A3 (both BD Biosciences). Data analysis was performed using the FlowJo software (Treestar, Ashland, OR). FACS sorting was performed with an AriaFusion or AriaIIIu (BD Biosciences).

### Quantitative RT-PCR

T cells were subjected to RNA extraction utilizing the RNeasy Micro Kit (QIAGEN). RNA from the renal cortex was isolated with the NucleoSpin Kit (Macharey-Nagel, Düren, Germany) in compliance with the manufacturer’s recommended protocol. Subsequently, the RNA underwent reverse transcription using the High-Capacity cDNA Reverse Transcription Kit (Thermo Fisher, Waltham, MA), and the StepOnePlus Real-Time PCR system (Thermo Fisher, Waltham, MA) was utilized for measurement. The Taqman primers used for human *IL17A*, *IFNG*, *CSF2*, *ZC3H12D*, *ATF4*, *18S rRNA,* and murine *Il17a* were procured from Life Technologies (Carlsbad, CA).

### *In-vitro* stimulation of cells

Human and murine T cells were classified based on the surface markers and FACS sorted before stimulation. Trm cells were characterized as CD45^+^ CD4^+^ CD44^+^ CD69^+^ CD62L^-^ ; Tem cells as CD45^+^ CD4^+^ CD44^+^ CD69^-^ CD62L^-^ ; Tcm cells as CD45^+^ CD4^+^ CD44^+^ CD69^-^ CD62L^+^ ; Naïve T cells as CD45^+^ CD4^+^ CD44^-^ CD69^-^ CD62L^+^. The cells were exposed to phorbol 12-myristate 13-acetate (PMA, 50 ng/ml, Sigma-Aldrich) and ionomycin (1 μM, Sigma-Aldrich) for T-cell activation. To detect cytokine production, brefeldin A (10 μg/ml, Sigma-Aldrich) was added to inhibit cytokine secretion from cells. After 3.5 hours, cytokine production was measured using intracellular staining and flow cytometry techniques. To inhibit de novo mRNA transcription, 20 µg/ml actinomycin D (A1410, Sigma-Aldrich) was added to the medium 10 minutes before T-cell stimulation. For *ex-vivo* stimulation to measure eIF2α levels, 1-2 × 10^6^ Trm cells were cultured in a volume of 300 μl of IMDM medium containing 10% FCS, streptomycin, and penicillin in a 96-well plate pre-coated with anti-CD3Ab (2 μg/ml, Biolegend 100360), along with the addition of 5 μg/ml anti-CD28 antibody (Biolegend 102122) for a duration of 2 days. To induce the ISR, T cells were incubated with 10 μM Sal003 (Sigma-Aldrich, S4451), 20 μM Sephin1 (Sigma-Aldrich, SML1356), 20 μM Raphin1 (Sigma-Aldrich, SML2562), or 500 μM NaAsO_2_ (Sigma-Aldrich, S7400).

### Puromycin incorporation assay

T cells were cultured in the presence or absence of PMA (50 ng/ml, Sigma-Aldrich) and ionomycin (1 μM, Sigma-Aldrich) for 1 hour. Puromycin (5 μg/ml, Sigma-Aldrich) was added in the last 10 minutes. Cells were washed and stained for surface markers and fixable dead-cell stain and then fixed and permeabilized using Foxp3/Transcription Factor Staining Buffer (eBioscience). Puromycin was detected using anti-puromycin-AF647 (MABE343 clone 12D10, Sigma-Aldrich).

### Polysome profiling for mRNA translation efficiency analysis

Polysome profiling and mRNA extraction were conducted as described previously (*27, 58*). Human Trm cells sorted from healthy human kidney tissue were mixed with a murine carrier cell line (NIH/3T3) because the number of human Trm cells was insufficient to show monosome peaks. We relied on monosome peaks from a murine cell line because monosome peaks were observed with the same kinetics in humans and mice. Cells were lysed in 250 μl cell lysis buffer (10 mM Tris-HCl (pH 7.4), 5 mM MgCl2, 100 mM KCl, 1% Triton X-100, 2 mM DTT, and 100 μg/ml cycloheximide). Cells were shear opened with a 26-gauge needle by passing the lysate 8 times through it. A quantity of 300 μL of the cell lysate was loaded onto a 5-ml sucrose gradient (60% to 15% sucrose) dissolved in polysome buffer (50 mM HEPES-KOH, pH 7.4, 5 mM MgCl_2_, 100 mM KCl, 2 mM cycloheximide, and 2 mM DTT) and separated by ultracentrifugation at 148,900xg (Ti55 rotor, Beckman) for 1.5 hours at 4°C. Polysome fractions and non-polysome fractions were collected separately. RNA was extracted from each fraction by adding 0.1 volume of 10% SDS and one volume of acidic phenol-chloroform (5:1, pH 4.5), incubated at 65°C for 5 min, and centrifuged at 21,000xg for 5 min at 4°C to separate different phases. Equal volumes of acid phenol-chloroform were added to the aqueous phase, separated by centrifugation, and supplemented with an equal volume of chloroform:isoamyl alcohol (24:1). Upon separation, the aqueous phase was supplemented with 0.1 volume 3 M NaOAc (pH 5.5) and an equal volume of isopropanol, precipitated at −20°C for 3 h. RNA was pelleted at 21,000xg at 4°C, and the dried pellets were resuspended in water. RNA was subjected to reverse transcription for RT-PCR analysis, and the original mRNA amount in polysome and non-polysome fractions was calculated by RT-PCR using Taqman primers specific to human mRNA. mRNA translation efficiency was defined as the amount of mRNA in polysome fraction divided by the total mRNA amount of non-polysome and polysome fractions.

### Single-cell RNA sequencing

To carry out scRNA-seq, single-cell suspensions were derived from human and mouse kidney samples. Cell hashing was implemented according to the manufactureŕs instructions (BioLegend). For CITE-seq, monoclonal antibodies with corresponding barcodes were applied to samples (Biolegend, San Diego, CA). FACS-sorted CD45^+^ cells or CD3^+^ T cells were subjected to droplet-based single-cell analysis and transcriptome library preparation using the Chromium SingleCell 5’ Reagent Kits v2 according to the manufactureŕs protocols (10x Genomics, Pleasanton, CA). The generated scRNA-seq libraries were subjected to sequencing on a NovaSeq6000 system (100 cycles) (Illumina).

### Alignment, quality control, and pre-processing of scRNA-seq data

Quality control and scRNA-seq pre-processing were carried out as previously described (*59*). In brief, the Cell Ranger software pipeline (version 5.0.1, 10x Genomics) was utilized to perform the demultiplexing of cellular barcodes and mapping of reads to the reference genome (refdata-cellranger-hg19-1.2.0 (human) or refdata-gex-mm10-2020-A (mouse). Seurat (version 4.0.2) demultiplexing function HTODemux was used to demultiplex the hash-tag samples. We filtered out the cells with <500 genes, > 6000 genes, or > 5% mitochondrial genes. For CITE-seq, a pseudo-reference genome was built with cellranger mkref function in Cell Ranger. CITE-seq raw data were aligned to this pseudo-reference genome using Cell Ranger function cellranger count. The antibody-derived tags (ADT) data were integrated to single cell RNA-seq data using Seurat. Further information is described previously (*13*).

### Dimensionality reduction, clustering, enrichment analysis, and scores

The Seurat package (version 4.0.2) was used to conduct unsupervised clustering analysis on scRNA-seq data (*60*). Briefly, gene counts for cells were normalized by library size and log-transformed. To reduce batch effects, we employed the integration method implemented in the latest Seurat version 4 (function FindIntegrationAnchors and IntegrateData, dims = 1:30). The integrated matrix was then scaled by the ScaleData function (default parameters). To reduce dimensionality, principal component analysis was performed on the scaled data (function RunPCA, npcs = 30). Thirty principal components were determined using the ElbowPlot function to compute the KNN graph based on the Euclidean distance (function FindNeighbors), which then generated cell clusters using the function FindClusters. Uniform Manifold Approximation and Projection (UMAP) was used to visualize clustering results. The top differential expressed genes in each cluster were found using the FindAllMarkers function (min.pct = 0.1) and running Wilcoxon rank sum tests. The differential expression between clusters or groups was calculated by the FindMarkers function (min.pct = 0.1), which also included Wilcoxon rank sum tests. For enrichment analysis including Gene Ontology, Reactome, Canonical pathway, and protein-protein interaction analyses, metascape (*61*) was used. To calculate scores (RPG score and ISR score) in scRNA-seq data, Seurat function AddModuleScore was used. For both scores, all RPGs (*62*) and ISR-associated genes (*19, 31–34*) listed in heatmaps (Supplementary Figure 5) were included.

### Statistics

Statistical analysis was performed using GraphPad Prism (La Jolla, CA). The results are shown as mean ± SEM when presented as a bar graph or as single data points with the mean in a scatter dot plot. Differences between the two individual groups were compared using a two-tailed t test. In the case of three or more groups, a one-way ANOVA with Tukey’s multiple comparisons test was used. The correlation coefficient r was calculated using a Pearson correlation, and the corresponding P value was based on a t-distribution test.

## ACKNOWLEDGMENTS

FACS sorting was performed at the UKE FACS sorting core facility. Single-cell RNA sequencing was performed at the UKE Single-Cell Core Facility. Graphical images were produced using Servier Medical Art images (https://smart.servier.com) and TogoTV (https://togotv.dbcls.jp).

## CONTRIBUTIONS

Conceptualization: N.A. and U.P. Methodology: N.A., P.G., H.J.P., N.S., A.P., A.K., H.W., and N.G. Formal analysis: N.A., C.F.K., and U.P. Flow cytometry: N.A., P.G., and H.J.P. scRNA sequencing data analysis: N.A., P.G., and Y.Z. Data analysis: N.A., P.G., A.K., and A.P. Polysome profiling: N.A., S.D., and Z.I. Renal histology: N.A. and U.P. Patient cohorts: J.E, J.H.R., P.G., R.D.,C.F.K., and U.P. Writing original draft: N.A. and U.P. Writing review and editing: N.A., H.-W.M., C.F.K., and U.P. Visualization: N.A., and U.P. Supervision: N.G., H.-W.M, C.F.K., and U.P. Funding acquisition: H.-W.M., C.F.K., and U.P.

## DATA AVAILABILITY

scRNA-seq data of T cells from human healthy kidney tissue and Trm cell-induced murine kidneys are available at FigShare: https://figshare.com/s/7912de1afc7fd5bbefd4. The scRNA-seq data of the healthy colon and lung were obtained from GSE157477 (*21*) and Cross-Tissue Immune Cell Atlas (https://www.tissueimmunecellatlas.org/#publication) (*22*), respectively. All other data needed to verify the study’s conclusions are contained in the manuscript or the Supplementary Materials.

