## Supplementary Figures for "The integrated stress response/eIF2a pathway controls cytokine production in tissue-resident memory CD4^+^ T cells"

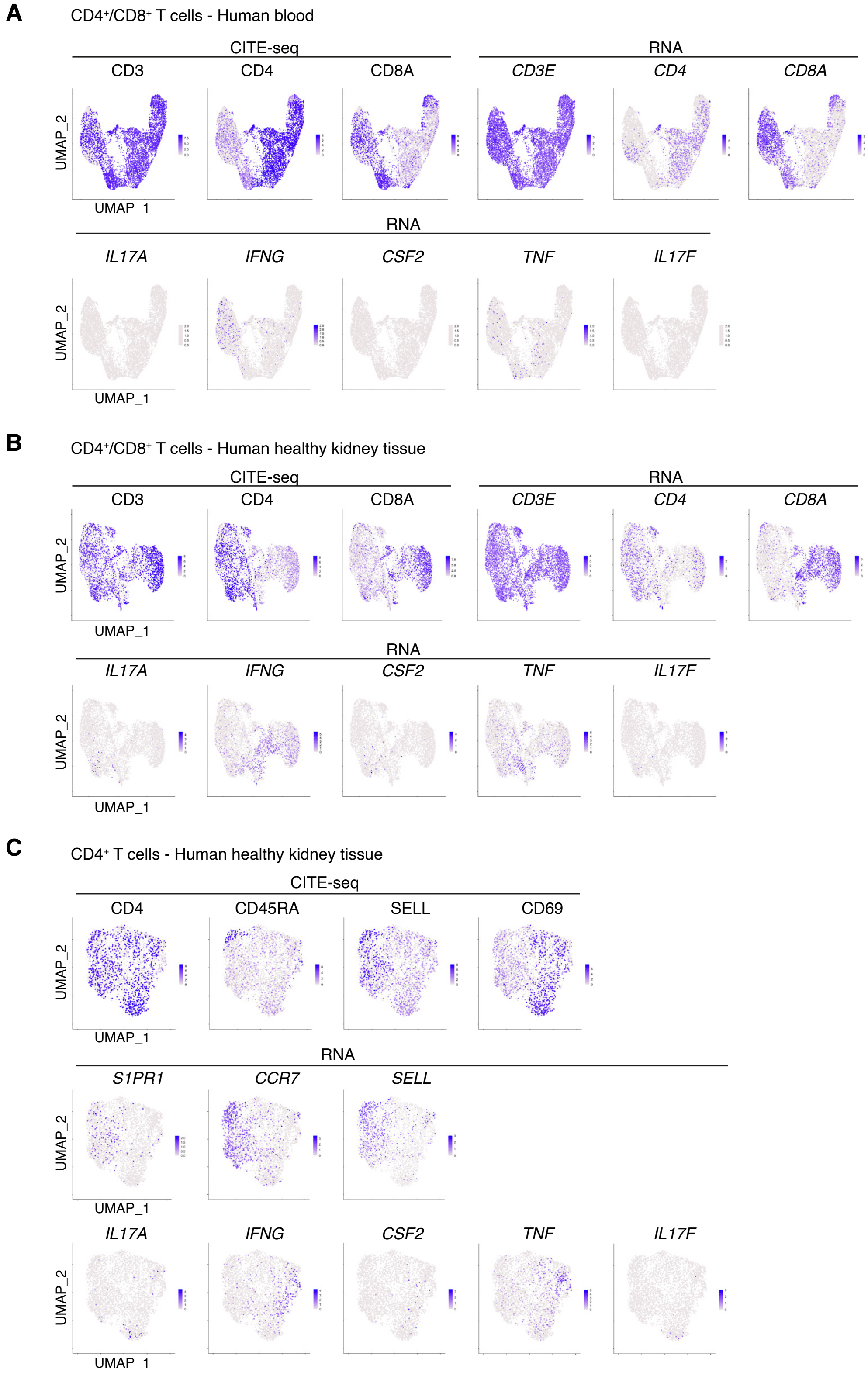


Supplementary Figure 1. scRNA-seq analysis of T cells from the human healthy kidney tissue and matched blood.
(A and B) UMAP plots showing marker protein (CITE-seq) and mRNA expression and cytokine mRNA expression of $CD4^{+}$ or $CD8^{+}$ T cells in the blood (A) and kidney (B).
(C) UMAP plots showing marker protein and mRNA expression and cytokine mRNA expression of $CD4^{+}$ T cells in the kidney.

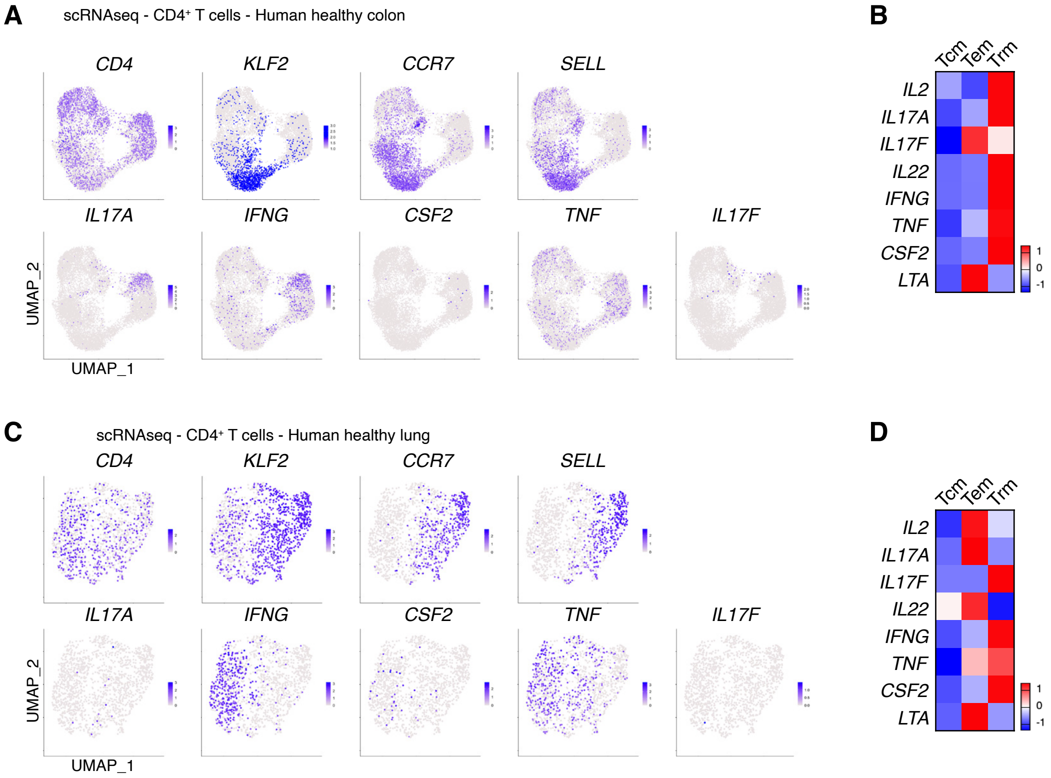


Supplementary Figure 2. Cytokine mRNA expression of the $CD4^+$ T cells from human healthy colon and lung.
(A and B) UMAP plots showing marker genes and cytokine mRNA expression (A) and a heatmap showing cytokine mRNA expression (B) in different $CD4^+$ memory T cell subsets from the heatlhy colon.
(C and D) UMAP plots showing marker genes and cytokine mRNA expression (C) and a heatmap showing cytokine mRNA expression (D) in different $CD4^+$ memory T cell subsets from the heatlhy lung.

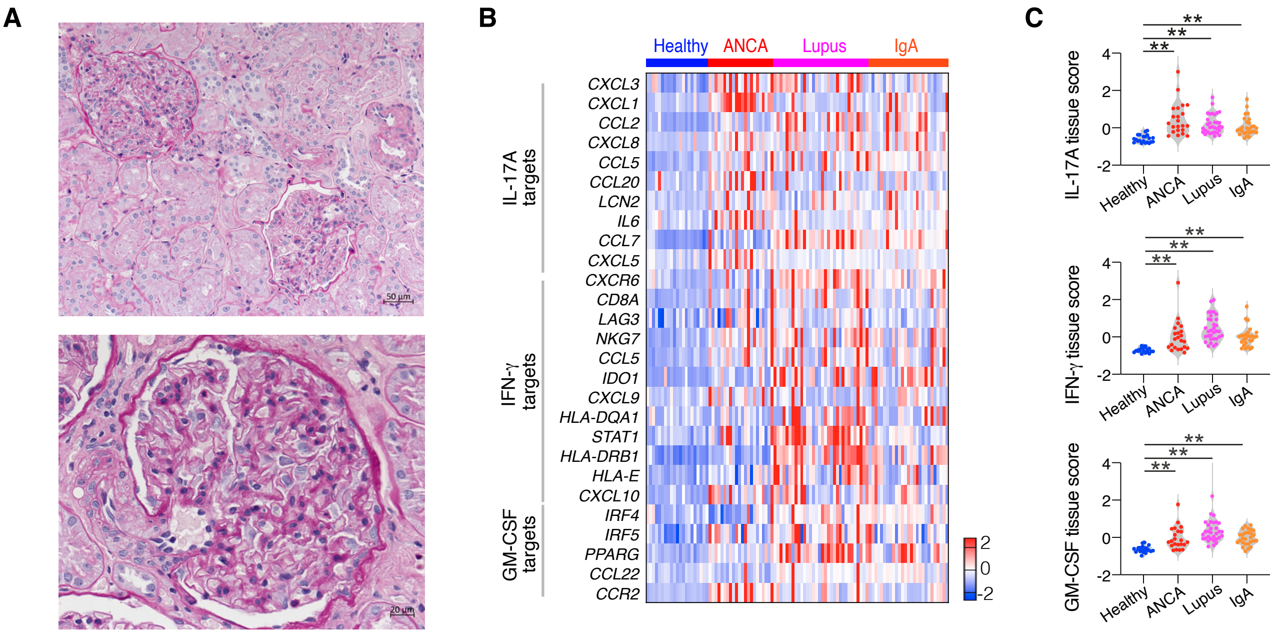


Supplementary Figure 3. Inflammatory response is not observed in healthy human kidney tissue.
(A) Representative PAS staing of human healthy kidney tissue.
(B and C) Heatmap showing the expression of cytokine-responsive genes in the kidney (B) and quantification of the gene expression as a tissue score (C). (* P<0.05, ** P<0.01)

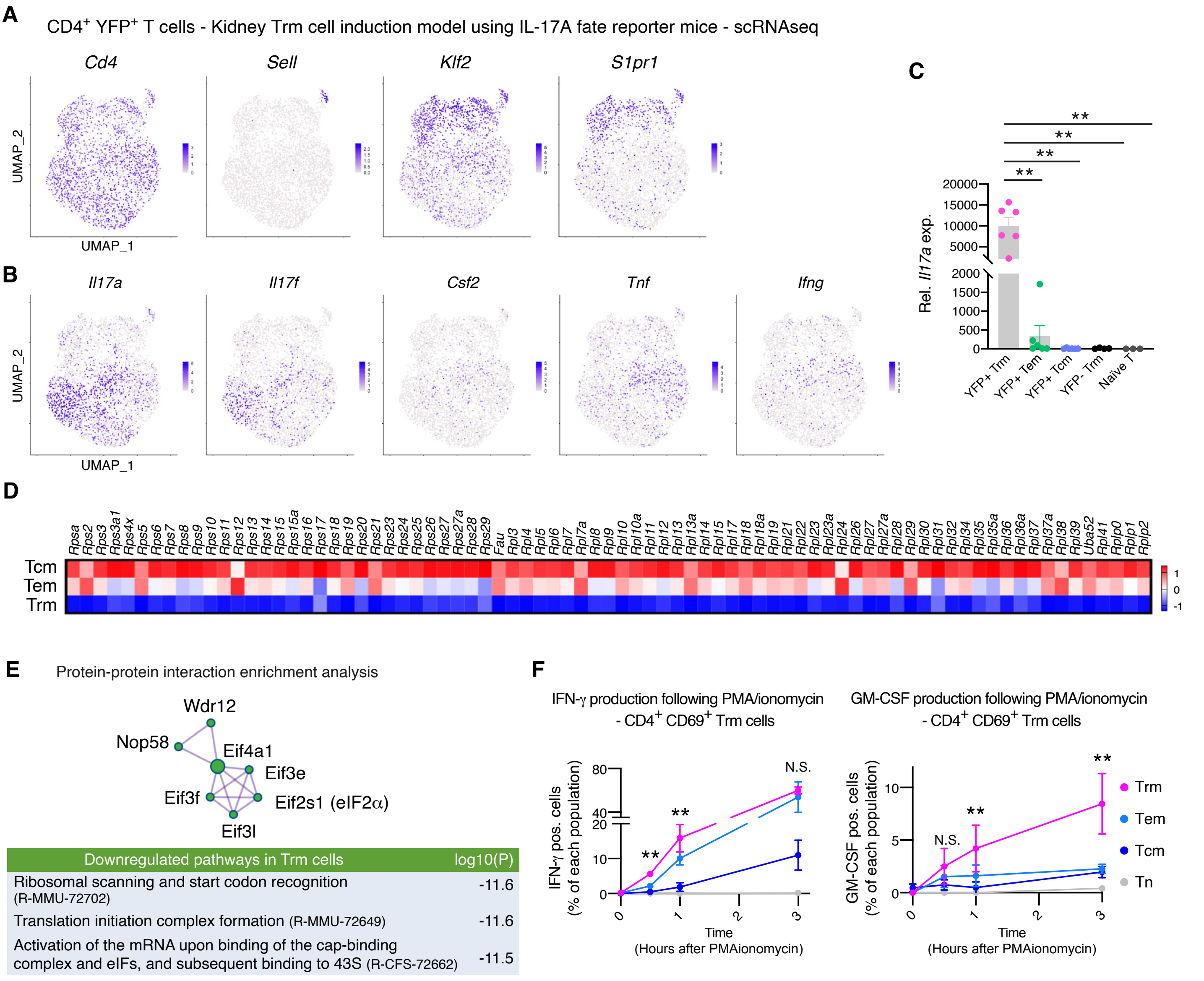


Supplementary Figure 4. Murine renal Trm cells express high level cytokine mRNA and produce cytokine protein immediately after activation.

Mice were infected with S. aureus, treated with ampicillin, and analyzed at least 2 months after initiating the administration of antibiotics.
(A and B) scRNA-seq analysis of CD4+ YFP+ T cells from the kidney. UMAP plots showing marker genes expression (A) and proinflammatory cytokine mRNA (B).
(C) Realtime RT-PCR analysis for *Il17a* mRNA expression in sorted renal T cells. Data are mean + S.E.M.
(D) Heatmap showing expression of mRNAs conding for ribosomal protein genes (scRNA-seq data).
(E) Protein-protein interaction enrichment analysis showing downregulated pathways in Trm cells (scRNA-seq data).
(F) Cytokine production by different renal T cell populations after PMA/ionomycin stimulation (n = 6 for each group).
(* $P < 0.05$, ** $P < 0.01$)

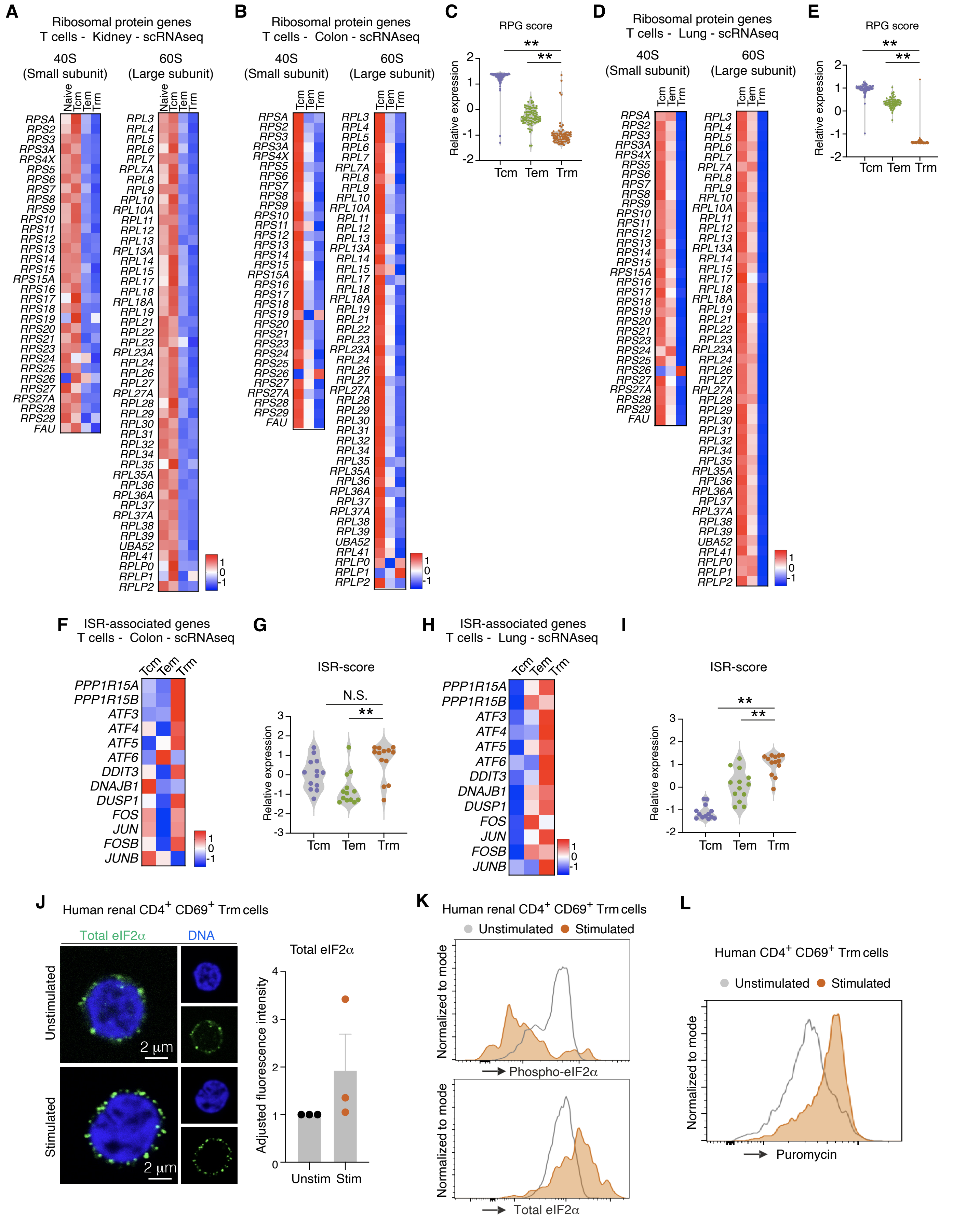


Supplementary Figure 5. Ribosomal protein genes (RPGs) and integrated stress response (ISR)-associated genes in T cells.
(A-E) RPGs expression shown in a heatmap and RPG score in different T cell subsets isolated from the human healthy kidney (A), colon (B and C) , and lung (D and E) tissue.
(F-I) Heatmap showing the expression of ISR-associated genes and ISR score in different T cell subsets in the human healthy colon (F and G) and lung (H and I).
(J) Representative images of total eIF2α staining in sorted Trm cells with or without PMA/ionomycin stimulation.
Data are mean + S.E.M.
(K) Representative histograms of phospho- and total eIF2α in Trm cells with or without CD3/CD28 Ab stimulation.
(L) Global protein synthesis was assessed with a puromycin uptake assay using Trm cells with or without PMA/ionomycin stimulation.
(* P<0.05, ** P<0.01).

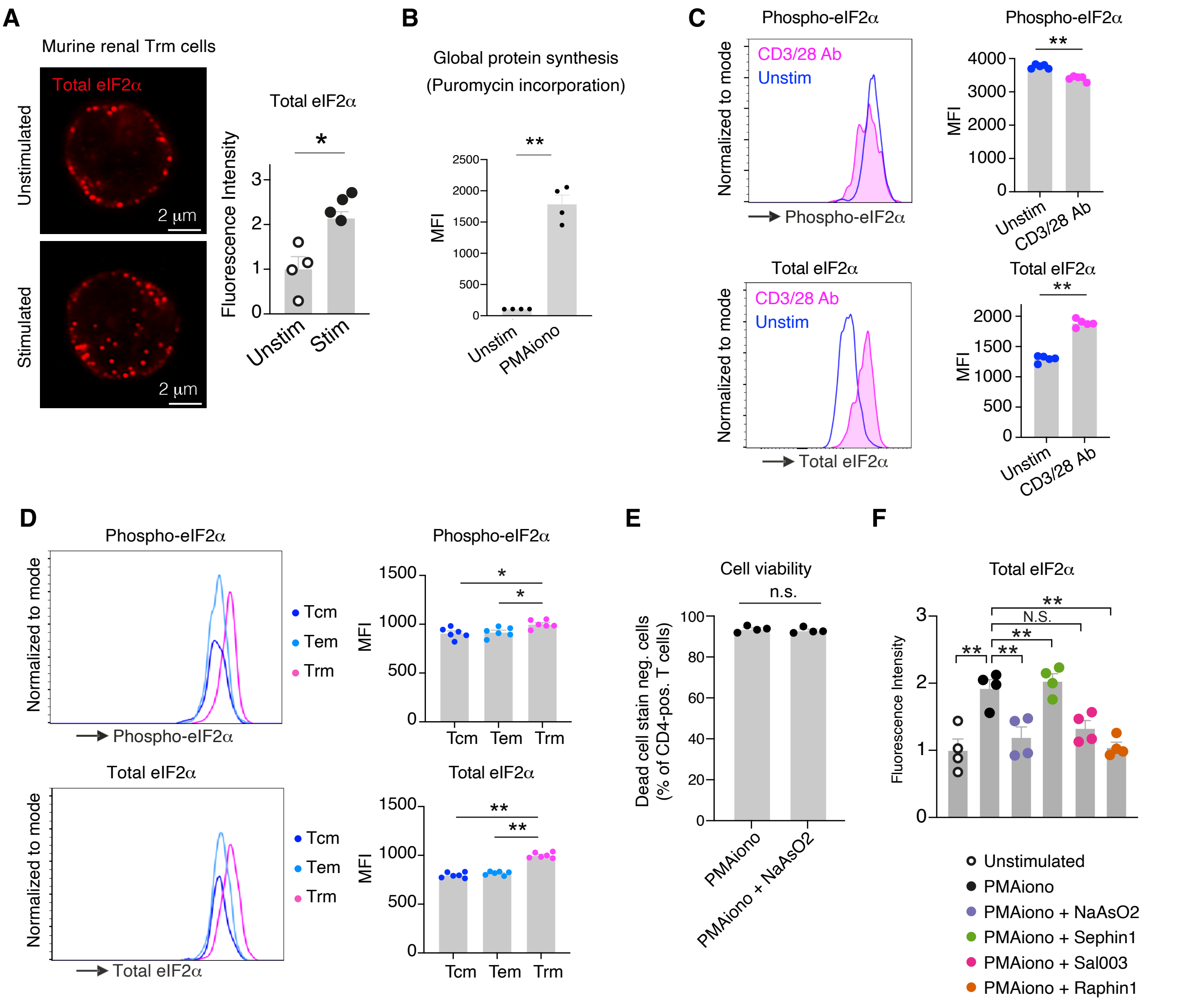


Supplementary Figure 6. Analysis of ISR in murine CD4+ CD69+ Trm cells.

(A) Representative images of total eIF2α staining in sorted Trm cells with or without PMA/ionomycin stimulation.

(B) Global protein synthesis of stimulated and unstimulated Trm cells analyzed by puromycin-uptake assay.

(C) Phospho- and total eIF2α levels in renal Trm cells stimulated with CD3/28Ab ex-vivo for 2 days.

(D) Phospho- and total eIF2α levels in different renal memory T cell subsets.

(E) Live cell percentage of Trm cells after treatment with PMAionomycin with or without $NaAsO_2$.

(F) Total eIF2α levels in Trm cells stimulated with PMA/ionomycin in the presence of indicated compounds (n = 4).

Data are mean + S.E.M. (* $P < 0.05$, ** $P < 0.01$)

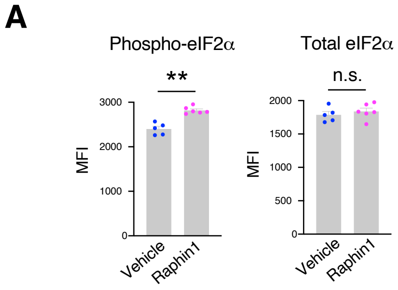


Supplementary Figure 7. Raphin1 increases phospho-eIF2α levels of Trm cells *in vivo*.
(A) eIF2α levels of renal Trm cells from nephritic mice 4 hours after Raphin1 application by oral gavage. eIF2α levels were measured by flow cytometry. Data are mean + S.E.M (* $P < 0.05$, ** $P < 0.01$).

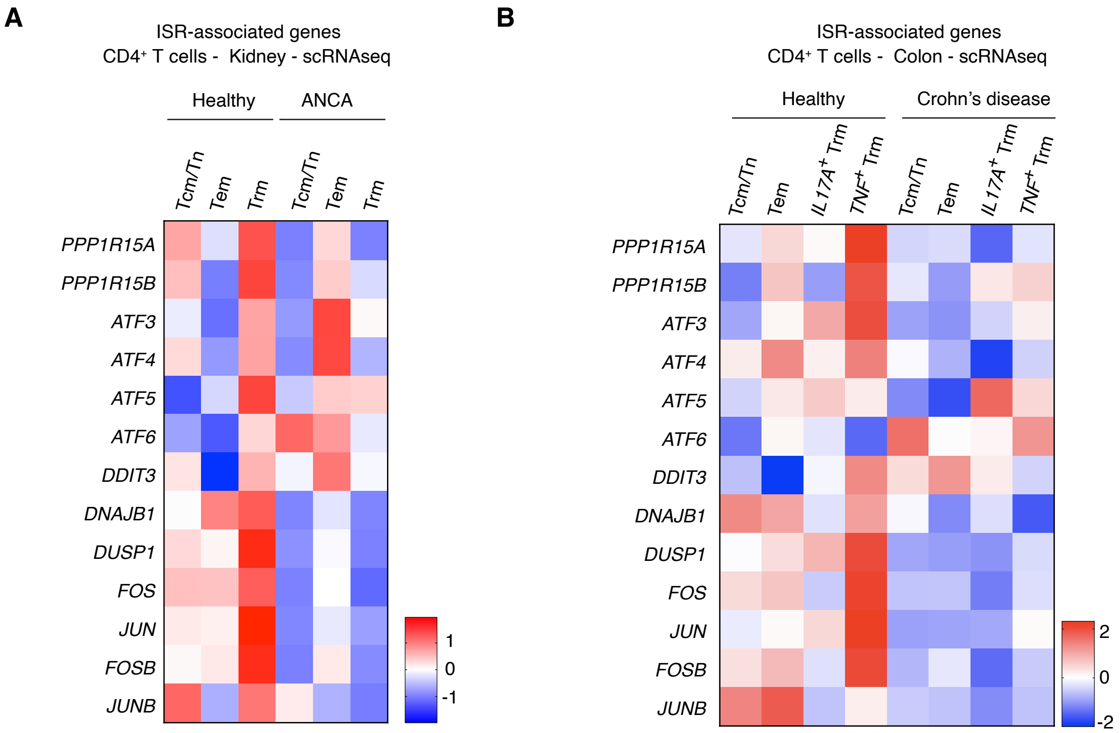


Supplementary Figure 8. scRNA-seq analysis of T cells from patients with ANCA-GN or Crohn's disease. (A and B) Heatmaps showing expression of ISR-associated genes in different T cell subsets isolated from patients with ANCA-GN (A) or Crohn's disease (B).

A

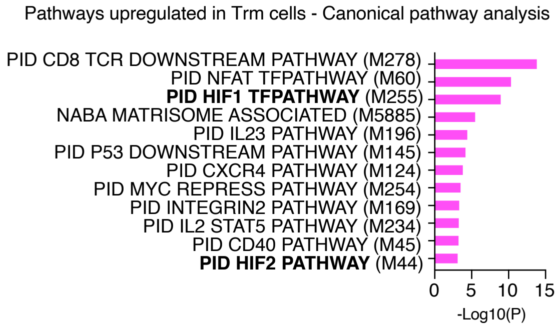


Supplementary Figure 9. Hypoxia-associated pathways are upregulated in human renal Trm cells.
(A) Pathways upregulated in Trm cells compared with non-Trm cells in canonical pathway analysis.
